# Tracing Homopolymers in *Oikopleura dioica*’s mitogenome

**DOI:** 10.1101/2024.05.16.594446

**Authors:** Nicolas Dierckxsens, Kosei Watanabe, Yongkai Tan, Aki Masunaga, Michael J. Mansfield, Jiashun Miao, Nicholas M. Luscombe, Charles Plessy

**Affiliations:** Genomics and Regulatory Systems Unit, Okinawa Institute of Science and Technology Graduate University, Okinawa, Japan; Keio University School of Medicine, Tokyo, Japan

**Keywords:** tunicate, *Oikopleura dioica*, homopolymers, larvacean, Appendicularia

## Abstract

*Oikopleura dioica* is a planktonic tunicate (Appendicularia class) found extensively across the marine waters of the globe. The genome of a single male individual collected from Okinawa, Japan was sequenced using the single-molecule PacBio Hi-Fi method and assembled with NOVOLoci. The mitogenome is 39,268 bp long, featuring a large control region of around 22,000 bp. We annotated the proteins atp6, cob, cox1, cox2, cox3, nad1, nad4 and nad5, and found one more open reading frame that did not match any known gene. This study marks the first complete mitogenome assembly for an appendicularian, and reveals that A and T homopolymers cumulatively account for nearly half of its length. This reference sequence will be an asset for environmental DNA and phylogenetic studies.

http://dx.doi.org/10.1364/ao.XX.XXXXXX

## 1. SIGNIFICANCE STATEMENT

Appendicularians, such as *Oikopleura dioica*, are planktonic tunicates which play an important role in the carbon cycle and the food web. Until this year, no annotated mitochondrial genome assembly was available for any appendicularian species. This lack of reference data hampers studies of their geographical ecology using environmental DNA sequencing (eDNA), which relies on the mitochondrial *cytochrome oxidase 1* sequence as a taxonomic barcode, resulting in appendicularians often being categorised as “uncultured eukaryotes” by default. Moreover, the mitochondrial transcriptome of *O. dioica* is subjected to post-transcriptional editing, which further complicates the prediction of DNA sequences from RNA and vice-versa. Therefore, curated reference annotation of appendicularian mitogenomes are essential not only for eDNA surveys, but also for supporting phylogenomic studies of the emergence of vertebrates, to which tunicates are the sister group, and for the enabling studies of the mechanism of homopolymer post-transcriptional edition. The complete mitochondrial genome assembly and its annotation published in this manuscript reveal the principles of *Oikopleura* mitogenome organisation and pave the way for the important studies summarised above.

## 2. INTRODUCTION

*Oikopleura dioica* Fol, 1872 [1], a marine tunicate within the Appendicularia class (larvacean), is notable for its widespread distribution across oceans. Throughout its lifecycle, *O. dioica* remains adrift, carried by ocean currents, and constructs a distinctive cellulose apparatus called the “house”, which it uses for protection and food collection. *O. dioica* frequently replaces its house when it becomes clogged, contributing to marine snow as discarded houses descend to the ocean floor. This process is important for Earth’s carbon cycling, emphasizing the species’ ecological significance [2].

We reported earlier the possibility of cryptic speciation in *O. dioica* [3], and that *cox1* nucleotide sequence similarity could be as low as ∼85 % similar when comparing *O. dioica* isolated from Europe [4][5], Okinawa [6], or the main Japanese islands [7][8]. Appropriate adjustment of the taxonomy is currently being discussed in the tunicate community. Nevertheless the evolutionary distance between these *O. dioica* cryptic species is large enough that separate reference sequences are needed to support applications of molecular biology to experimental research and taxonomic surveys using mitochondrial sequences as barcodes. This evolutionary distance also provides useful data to support phylogenetic studies of appendicularians and tunicates, which are needed to understand the the evolution of early vertebrates.

The mitogenome of *O. dioica* was first characterized in the supplemental material of Denoeud *et al*. [4] in 2010, revealing 8 genes (*cox1, cox2, cox3, nad1, nad4, nad5, cob*, and *atp6* ). The existence of *nad2* remained under question. The ascidian mitochondrial genetic code was used by Denoeud *et al*. to translate these coding genes. Furthermore, in 2019 we used multiple sequence alignments of Cox1 and Cob to demonstrate that this genetic code arose early in the tunicate history and confirm its appropriateness for *O. dioica* sequences [9] despite they are not ascidians.

Denoeud *et al*. also noted the presence of poly-T insertions in coding regions, and stated the hypothesis that they are reduced to 6-mers in the transcriptome with a RNA editing mechanism. In 2021, we produced a partial assembly of 9,225 kbp containing the previously reported genes except *nad5* [6]. Unfortunately the length of the the poly-T insertions could not be assessed with confidence because of limitations in the Nanopore basecalling technology, and the use of post-assembly polishing methods. Neither of the two studies could confirm whether the *O. dioica* genome was a single circle, linear, or a collection of minicircles like in *Salpa thompsoni* [10]. Sequencing complete *O. dioica* mitochondrial genomes has remained a challenge until now because of the lack of technologies accurate over long homopolymers, and the lack of software capable to assemble these regions.

## 3. MATERIALS AND METHODS

### A. Sample collection and DNA extraction

We collected *Oikopleura dioica* specimens at Ishikawa harbor, Okinawa, Japan (26.114N, 127.665E) in June 2018. The samples were washed with 5 mL filtered autoclaved seawater three times, and then resuspended in 200 µL lysis buffer from the MagAttract HMW DNA Kit (Qiagen, 67563) with 20 µL of 10 µg/mL proteinase K and incubated for 1 h at 56℃ After adding 50 µL of 5 M NaCl, the mixture was centrifuged at 5000 × g at 4℃ for 15 min. The supernatant was transferred into a new microtube with 400 µL of 100% EtOH and 5 µL of glycogen (20 mg/mL) and cooled at *−*80℃ for 20 min. After centrifuging at 6250 × g, 4℃ for 5 min, the supernatant was removed. The pellet was washed with 1 mL of cold 70% ethanol, centrifuged, and air-dried 5 min. Finally, the DNA was resuspended in molecular biology grade H2O for quantitation using a Qubit 3 Fluorometer (Thermo Fisher Scientific, Q32850), and quality-controlled using an Agilent 4200 TapeStation (Agilent, 5067–5365).

### B. Sequencing and Assembly

The genome of a single individual (“I25”) was sequenced on PacBio Sequel II using a low-input HiFi library kit. Genomic DNA was sheared with Megarupter3 to an average size of 10 kbp. Library size distribution and concentration were assessed using the FEMTO Pulse system.

The sequence reads were assembled with NOVOLoci [11], a newly developed targeted haplotype-aware assembler based on the same principle as the organelle assembler NOVOPlasty [12], using a seed sequence from the *O. dioica* OKI2018_I69_1.0 sequence [13].

### C. Annotation

We attempted to annotate the assembly with MITOS [14] using the ascidian mitochondrial genetic code [9]. However, the long length of the homopolymers (Fig.1, Fig.2, and Table 1) made it problematic for MITOS to detect entire genes with enough precision and lead to a large number of false positives. We hence compared the genome to the Trinity transcriptome assembly of OKI2018_I69 [13] to detect each gene. We queried the transcript models using amino acid sequences from MITOS or from related tunicates using `tblastn` [15]. The matched models, which were polycistronic, are shown in Table 2 and provided in the supplementary material. We determined the extent of each gene’s coding sequence using the `getorf` command from EMBOSS [16] with `-table 13`. The region encoding the gene was then aligned to the genome to discover the locations of poly-A or poly-T insertions, by Smith-Waterman alignments with the `water` command.

**Fig. 1.**
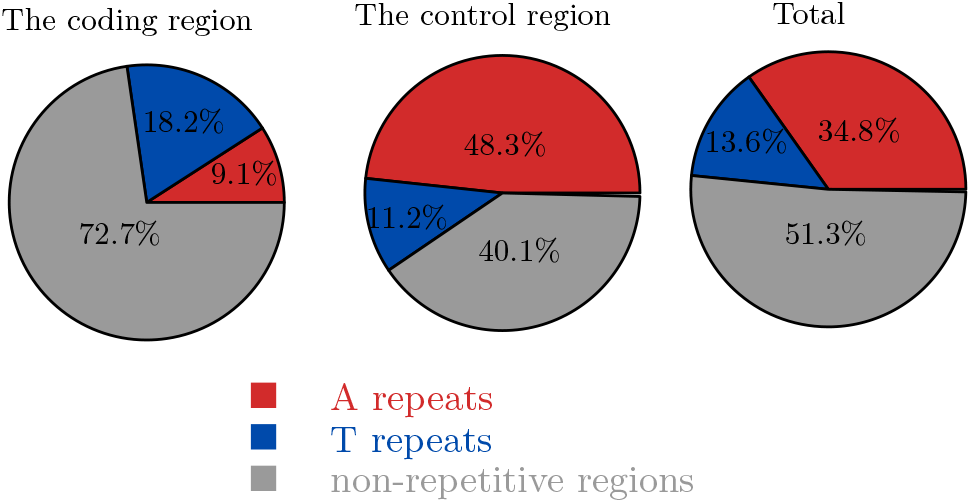
Pie charts showing he proportion of the homopolymers of 6 successive bases or longer. C repeats do not exist and G repeats are slight in proportion, thus excluded from the charts.

**Fig. 2.**
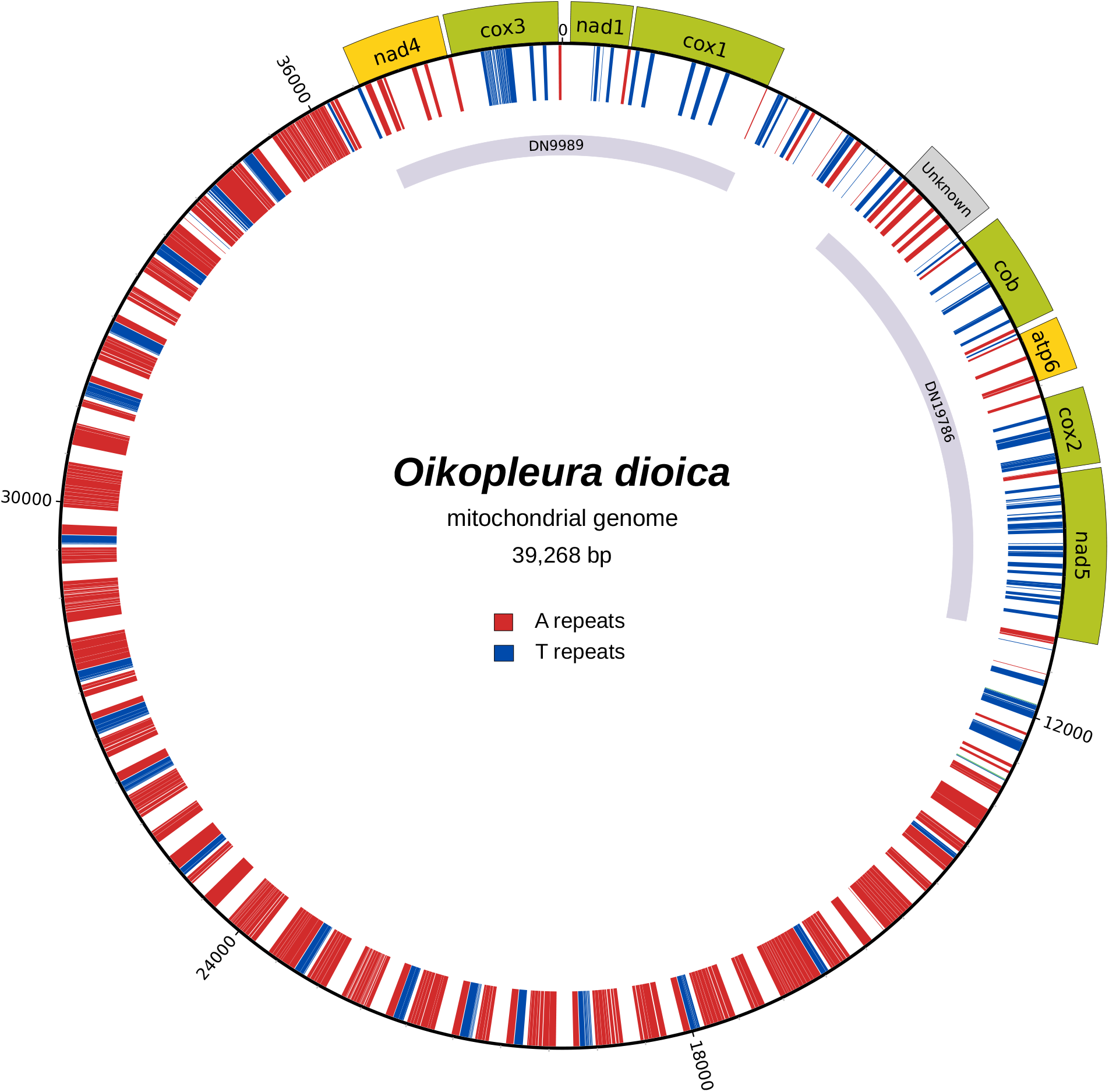
Circular plot drawn by Circos version 0.69-9 [20]. Known protein-coding gene symbols are displayed on a green background for plus-strand genes and yellow for minus strand. Unknown proteins and transcript models are displayed in gray. The inner circle illustrates the repetitive regions; 6 or more successive A or T’s are coloured respectively red and blue.

**Table 1.** Coordinates of the annotated genes, IDs of transcripts used for the annotation, number of edited homopolymers.

| gene | start | end | number of n-mers | strand | transcript ID |
| --- | --- | --- | --- | --- | --- |
| *nd1* | 98 | 820 | 4 | + | TRINITY_DN9989_c0_g1_i1 |
| *cox1* | 859 | 2625 | 5 | + | TRINITY_DN9989_c0_g1_i1 |
| *cob* | 5787 | 7021 | 7 | + | TRINITY_DN19786_c0_g1_i4 |
| *cox2* | 7963 | 8857 | 4 | + | TRINITY_DN19786_c0_g1_i4 |
| *nd5* | 8914 | 10959 | 12 | + | TRINITY_DN19786_c0_g1_i4 |
| *cox3* | 37885 | 39219 | 3 | + | TRINITY_DN9989_c0_g1_i1 |
| *atp6* | 7110 | 7738 | 4 | - | TRINITY_DN19786_c0_g1_i4 |
| *nd4* | 36676 | 37828 | 5 | - | TRINITY_DN9989_c0_g1_i1 |

### D. Phylogenetic tree

We collected chordate *cox1* nulceotide sequences from publicly available complete mitogenomes assemblies. In addition, we also extracted *cox1* from genomic scaffolds for *M. erythrocephalus* (SCLF01725989), *B. stygius* (SCLE01415711) and *O. longicauda* (SCLD01101138) [17]. *O. dioica* sequences are from this work (I25) or were extracted from transcriptomes [5][7][6]. A copy of the sequences is in the supplementary material. We aligned the codons within sequences with Clustal Omega [18] and Seaview [19], and computed a tree with IQTREE with parameters `–polytomy –ufboot 1000 -m MFP -au -zb 1000`.

## 4. RESULTS

We assembled a circular mitogenome of length 39,283 bp, containing 39.7% A homopolymers of length 6 or more and 13.6% T homopolymers of length 6 or more (Fig. 1). While existing algorithms were incapable of assembling the complete mitochondrial genome, NOVOLoci succeeded by step-wise extending the seed into a complete circular genome. We confirmed the existence of each gene reported earlier [4] (Fig. 1, Table 1). All coding genes ended with TAA juxtaposing A-homopolymers of length greater than 6. No A-homopolymer was found within open reading frames (a feature which in retrospect makes the genome very easy to annotate), with the possible exception of *cob*, for which we lack phylogenomic or proteomic evidence to determine if the translation starts before or after an A-homopolymer present at the beginning of the open reading frame. Every T-homopolymer longer than 6 in the coding genes was reduced to 6-mers in the transcriptome. We note that some poly-T regions had non-T insertions which were also removed. Visual inspection of sequence read alignments to the genome confirmed the accuracy of the sequence at these insertions.

The region between *cox1* and *cob* does not encode known proteins, but we found an long open reading frame at the same position where Denoeud *et al*. hypothesised *nad2*. Unfortunately, the identity of this gene could not deciphered as its sequence did not have matches in the NCBI BLAST databases. The other open reading frames of the region were all shorter and also without known protein or nucleotide matches. The *cox1* sequence reported here is 99.4 % similar to the transcript model we used as a seed, and encodes for the same protein sequence.

To place our reference genome into context and to illustrate appendicularian relationship to other taxa, we computed a phylogenetic tree on 87 codon-aligned *cox1* chordate sequences (Fig. 3). The appendicularian branch placement is different from our previous analysis based on nuclear gene orthogroups[22], but the discrepancy may be caused by long branch attraction. We also note that the *Ciona* genus grouped with *Aplousobranchia* instead of the *Phlebobranchia* where current taxonomy places it, but this discrepancy has been observed in numerous phylogenomic analysis before. Finally, the division between *O. dioica* and other appendicularians corresponds to the *Coecaria* (*O. longicauda* group) and *Vexillaria* (*O. dioica* group) and is well supported by taxonomy [23] and genomics [17]. Sequence identity between *O. dioica* and *O. longicauda* is below 60 %; no full-length *cox1* sequence is available for closer relatives of *O. dioica* such as *O. albicans* or *O. vanhoeffeni* [17].

**Fig. 3.**
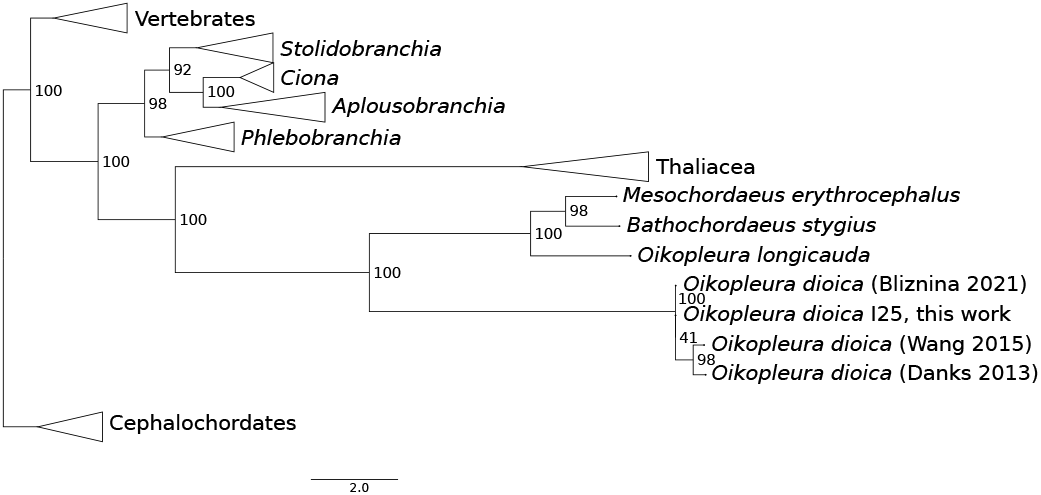
Relation between *cox1* sequenced this work and the other publicly available appendicularian sequences.

## 5. DISCUSSION AND CONCLUSION

We present here the first complete mitogenome of *Oikopleura dioica*. The assembly is circular, but we can not exclude the possibility that the genome is actually a linear concatenate or present in multiple copies in a circle, due to the length of the control region. Our findings confirm the presence of genes previously reported and identify an unusual high amount of poly-T insertions in the control region. Our annotation reveals a general principle that will facilitate the study of other mitogenomes containing homopolymer insertions in the *Oikopleura* genus. First, genes are separated by poly-A regions directly encoded in the genome. Second, poly-T insertions containing a few non-T bases may also be edited. The discovery of this principle enables the annotation of homopolymer-containing *Oikopleura* mitogenome in the absence of transcriptome annotation.

Our protein-coding gene annotation does not resolve the possible the loss of the ATP synthase subunit *atp8* and the dehydrogenase subunits *nad2, nad3, nad4L* and *nad6* since the last common ancestor with ascidians. Investigation of the ORFs between *cox1* and *cob* is currently hampered by the lack of sequence homology with other tunicates.

During the preparation of this manuscript, we became aware of the publication of a mitogenome assembly for an European isolate of *O. dioica* [21]. In order to avoid confusion caused by the high nucleotide divergence, we assigned our assembly to NCBI’s taxon ID 3071372 as “unclassified Oikopleura”, pending the needed taxonomic adjustments. These two assemblies will be an asset for detecting *O. dioica* in environmental DNA (eDNA) studies of diverse geographical regions, and open the way for comparative approaches to elucidate non-coding regions and small ORFs. Future production of mitogenomes from the *Oikopleura* genus will be needed will increase the power of these methods.

We confirmed that the gene order in our assembly is identical to that of Denoeud *et al*., 2010 [4], extracted from a Norwegian laboratory line, and that of Bliznina *et al*., 2021 [6], extracted from our laboratory line from Okinawa, Japan. We recently published that in the nuclear genome, the order of protein-coding genes is “scrambled” when comparing these two *O. dioica* lines [22]. It is therefore noticeable that although the order of genes is said to be less conserved in mitogenomes [24], no change took place in *O. dioica* at the time scale separating these two populations.

The mechanism of poly-T editing in the mitochondrial mRNAs is not understood. Despite the existence of this editing is common knowledge in the scientific community working with appendicularian mitochondrial sequences, our publication and that of [21] are only the second ones after Denoeud *et al*. [4] to provide evidence matching genome and transcriptome data. Because of 1) this reason, 2) the uncertainty on the coding gene count, and 3) the absence of ncRNA annotation, we were only allowed to deposit the genome’s sequence in GenBank as “UNVERIFIED”, with no annotation, which we provide as supplemental material. This is unfortunate because it hides essential taxonomic information to metagenomics and environmental DNA studies that rely on the existence of annotated hits with strong sequence similarity in public databases. This peer-reviewed publication aims as building up the evidence available to curators regarding the existence of homopolymer editing in the *Oikopleura* genus, and the need to adjust database infrastructure so that this biological phenomenon can be properly representation in annotations distributed by public databases.

## 6. APPENDICES

Additional file: Supplemental data (10.5281/zenodo.11142978).

## 7. ETHICAL APPROVAL

No ethical issues were involved in this study.

## 8. AUTHOR CONTRIBUTIONS

AM collected samples, YT cultured and sequenced them. ND, KW, MJM, JM and CP performed bioinformatics analysis. KW and ND drafted the manuscript. CP, and NML critically revised the manuscript. All authors approved the final manuscript and agreed to be accountable for all aspects of this work.

## 10. DATA AVAILABILITY

The mitochondrial genome sequence was deposited in GenBank under the accession number PP146516. The annotation and the sequences used to compute Figure 3’s tree are available in Zenodo under DOI https://doi.org/10.5281/zenodo.11142978.

## 11. ACKNOWLEDGEMENTS

We thank the DNA Sequencing Section and the Scientific Computing and Data Analysis Section of the Research Support Division at OIST for their support. This work was supported by OIST core funding. M.J.M. acknowledges funding from the Japan Society for the Promotion of Science as a JSPS International Research Fellow (Luscombe Unit, Okinawa Institute of Science and Technology Graduate University).

